# Silent brains isolate the heartbeat-evoked potential

**DOI:** 10.64898/2026.07.22.740079

**Authors:** Rohan Kandasamy, Rania-Iman Virjee, David Carmichael, Sarah N. Garfinkel, Mahinda Yogarajah

## Abstract

**Introduction:** The heartbeat-evoked potential (HEP) is widely interpreted as an EEG/MEG marker of cortical processing of cardiac afferent signals. A central methodological challenge in HEP research is separating genuine cortical responses to cardiac afferent signals from heartbeat-related cardiac field artefact (CA). Because both neural HEP activity and cardiac artefact are time-locked to the same heartbeat, conventional averaging and blind-source correction methods lack an observable cardiac-only reference, making their separation difficult to validate.

**Methods:** Individuals with isoelectric-appearing cerebral activity i.e. those with significant brain injury leading to a lack of detectable brain activity at the scalp, have an absence of recordable HEP signal yet preserved cardiac field artifact in brain. We used clinically acquired EEGs with isoelectric-appearing cerebral activity and preserved cardiac activity as a negative control for non-neural heartbeat-locked scalp signal. Thirty-nine isoelectric participants were compared with 118 heart-rate-matched participants whose clinical EEGs were reported as normal. Participant-level heartbeat averages were analysed using covariate-adjusted spatiotemporal permutation tests. We characterised interindividual cardiac-artefact morphology and tested the primary group contrast across average-reference, surface-Laplacian and cardiac-artefact-directed ICA analyses, with pseudotrial correction for heartbeat-independent EEG activity.

**Results:** Cardiac-artefact morphology varied markedly across isoelectric participants, was predominantly posterior or posterolateral, and was not predicted by ECG morphology. We attempted delineation of the HEP from the CA by comparing the two groups, revealing a negative frontocentral cluster that persisted in control participants at 90–365 ms, p=0.16 and a posterior cluster at 145–310 ms, p=0.03. The topography of the first cluster aligns with the spatiotemporal characteristics of the early window of the HEP as identified in the literature. This cluster was negative in polarity at the scalp. After independent component analysis, the same comparison produced a less prominent cluster. There was no significant group difference in ECG.

**Conclusion:** These findings provide an important first empirical scalp-level dissociation of the early HEP from CA using an inversion-of-ground-truth approach. A cluster of difference consistent with typical distributions of the early HEP, suggests that this part of the HEP is a distinct, frontocentral, negative cortical response. The effects of conventional independent component analysis for removing the CA suggest a risk of also removing veridical HEP signal with standard correction methods.

**SIGNIFICANCE STATEMENT:** This study uses an inversion of ground truth approach to characterise the cardiac artefact seen in EEG in the absence of brain activity, and thereby better characterise the heartbeat-evoked potential. This is of benefit for future interoceptive research as characterisation of the artefact can future approaches to preprocessing, and knowledge of the polarity of an evoked potential allows inference about whether a change in the amplitude represents more or less event-related activity.

**HIGHLIGHTS:**

1. Heartbeat-related artefact varies in distribution between individuals.
2. The heartbeat evoked potential is distinct from the cardiac artefact.
3. The early part of the heartbeat evoked potential is negative.

## INTRODUCTION

Despite four decades of research into the heartbeat evoked potential (HEP)(1), which is the evoked potential time-locked to cardiac activity, HEP responses have been solely captured entangled with cardiac artifact. EEG (scalp(1) or intracranial(2, 3)) or MEG(4) has been shown to capture a central response to each heartbeat (the HEP). At the same time, the heart produces a cardiac field artefact (CFA, from the volume conduction of the heart’s electrical behaviour) and the cardioballistic artefact (CBA, movement artefact produced by the pulse)(5). The CBA and CFA may be orders of magnitude larger than the HEP(6) (sometimes sufficient to be visible, unaveraged, to the naked eye).

Averaging cannot separate signals with the same temporal trigger, and conventional blind-source correction methods have no external criterion for determining whether a cardiac component also contains HEP. The field therefore lacks an empirical reference for the heartbeat-locked signal that would remain in the absence of detectable cerebral activity. An effect of their never having been an analysis of the HEP without a known baseline has meant the polarity (and the morphology) of the HEP is unknown.

A reproducible evoked component has a characteristic source configuration. That source configuration produces a scalp field with a particular polarity under a specified montage. Once the component’s polarity is established, changes along that field can be interpreted as greater or lesser expression of that component. For the HEP, this interpretation has been obstructed because the apparent zero or baseline is contaminated by the CA(6), to an unknown extent.

An growing body of research applies the HEP in the context of clinical neuroscience(7), social neuroscience(8), computational neuroscience(9). The HEP has been shown to be modulated by both trait(10, 11) level and state(12–14) level differences, for example amplitude modulation attributable to changes in direction of attention (towards and away from the body) or relating to the emotionality of presented faces(14). Better quantifying the HEP, though, has the potential to enhance the veracity of HEP, particularly interindividual trait differences.

We exploit a negative control: clinical EEG recordings with isoelectric-appearing cerebral activity but ongoing cardiac activity. These were taken from individuals who have a significant brain injury leading to a lack of detectable brain activity at the scalp. In these recordings, the heartbeat-locked scalp signal should be dominated by non-neural cardiac artefact. Comparing them with identically processed, heart-rate-matched clinical EEGs containing ordinary cerebral activity provides an empirical test of where and when the latter depart from the cardiac-artefact distribution. Any residual HEP in the isoelectric group, with the same scalp field, would attenuate the group contrast and therefore make the test conservative. The principal alternative explanation is systematic difference in cardiac artefact between groups.

We therefore had three aims: how variable is heartbeat-locked cardiac artefact during isoelectric EEG; where and when does heartbeat-locked EEG in controls depart from this negative control (and what is its polarity); and does that difference remain under artefact-removal procedures?

## METHODS AND MATERIALS

### Ethics Statement

This study was performed using a published dataset under the aegis of the Temple University EEG Dataset’s own ethical approvals(15).

### EEG selection

The data was selected from the Temple University EEG dataset (TUEG)(15), categorising metadata of which had been extracted from an older version of the corpus for another study(16). The EEG was then visually inspected by a clinical EEG reviewer (RK) to confirm isoelectricity, visually inspected with a sensitivity of 5 microvolts per centimetre, assuring that the EEG met ACNS guidelines for brain death(17), with confirmatory computational methods discussed later. Controls which had been expertly reported as normal (in the reports in the TUEG dataset) were selected to match the heart rates of the IE group. This BPM-matched (BM) group were matched with 3 normal EEG records for each in the IE group, with a tolerance of 2 beats per minute, with the expectation that some would be excluded due to artefact. Due to the nature of the dataset, detailed clinical information was not available, beyond that already extracted from the EEG reports.

Records were 10-20 clinical montage studies, occasionally with added leads. The analysis was limited to the shared standard montage.

### Preprocessing

EEG handling was supported using the MNE-Python(18), using the HEPPy pipeline, which was updated to allow more flexible analysis for this project. The identified records were reformatted, where necessary, to an average reference. This was chosen because mastoids may produce a nonuniform (and perhaps enhanced) cardiac field.

EEG records were preprocessed first with pyprep(19), and filtered between 0.5 and 100Hz. At this stage, if there were too few leads to perform reliable interpolation, the participant was excluded. According to standard pyprep methods this was set as 25% of the available leads in the montage(19). ICA was performed, using the infomax algorithm and then artefactual components identified with the MNE-Iclabel(20, 21) function. This was used to identify and exclude components identified as anything other than “brain,” “heart” or “other”. We selected artefacts for exclusion to minimise the risk of modification of the HEP (and CA).

Once the data was pre-processed, the ECG lead was cleaned and analysed using neurokit2(22), and R-waves were labelled. Neurokit2’s inbuilt ECG preprocessing was used to clean the data, with its “promac” algorithm for peak identification. If too few (less than 30 within a single record) were identified, the record was excluded at this step. 30 was chosen because a total of 100 beats were required, and few participants would have more than 2 records. Including very asymmetrical records (for example one with 90 beats and one with 10) raised the concern that the beats in the shorter record would be more artefactual and ECG quality would be less reliable as it is based on within-record statistics. Epochs were then generated based on the identified R peaks. Epochs were excluded if the amplitude in the epoch exceeded 150 microvolts, emulating typical analytical pipelines(23), in the BM group. In the IE group, epochs were excluded with amplitudes over 10 microvolts, after windowing to exclude the QRS complexes, as any activity was definitionally artefactual, but 10 microvolts was sufficiently high a value to not exclude the T wave. This had the added benefit of adding an objective, automated layer to the human inspection for isoelectricity.

Isoelectric EEG is not conceptually identical to brain death, despite being used for the diagnosis thereof, but the definition of isoelectricity is more pertinent for this study. We aim to exclude cortically-derived fields at the scalp. It requires that the scalp-detectable cortical HEP is absent in the isoelectric recordings, not that every neural structure is electrically silent. Any residual HEP with the same scalp field would attenuate the group contrast and therefore make the test conservative. The principal alternative explanation is systematic difference in cardiac artefact between groups. Candia-Rivera and Machado previously reported a single HEP analysis in a brain-dead patient(24), but as their focus was the characterisation of the HEP (and its absence) they used ICA to attempt to remove the CA. In their study, the HEP appeared absent, supporting the present study’s assumptions, however their methods did not permit a comparison between the CA and the HEP.

Epochs were centred on the R-peak (t=0). EEG was resampled to a common sampling rate of 200 Hz to ensure consistent time-grids across participants (as the recording frequencies varied between different records). Where participants had multiple records, these were concatenated after preprocessing to avoid pseudoreplication, using strict channel matching to ensure consistency in montages between recordings. Participants were excluded from the analysis if they had fewer than 100 usable epochs across all concatenated epochs.

Finally, a second ICA was performed with manual identification of components that resembled CA. To prevent bias in selection, the order of the components was randomised between records (across both groups), meaning that the components were assessed (by RK) without knowledge of whether they came from a BM or IE EEG. CA-like components were then removed from the EEG. For the components identified as CA, the relative amplitudes of the QRS-complex period and the post-T period of the cardiac epoch were recorded. The latter giving an estimate of the amplitude of the component outside the ECG-locked activity.

### Analyses

#### Morphological characteristics

We first characterised the morphology and interindividual variability of cardiac artefact in the isoelectric group. This dataset provides a rare chance to explore the typical scalp artefact coverage in (near) isolation. Their heartbeat-locked scalp signal was treated as an estimate of cardiac artefact in the absence of a scalp-detectable HEP.

The analysis was restricted to 100–900 ms after the R peak, corresponding to the interval in which cardiac artefact overlaps most closely with commonly studied HEP windows. The QRS complex itself was excluded because its substantially greater amplitude would otherwise dominate the clustering solution. The clustering was performed between the averaged cardiac artefacts for each participant, rather than individual epochs.

First, the data was time-warped (using linear interpolation) to align all epochs’ RR intervals to an arbitrary fixed duration of one unit (one second in the native time scale, N.B. this was solely for the clustering analysis).

We grouped participants by similarity of the morphology of their cardiac artefact, using spectral clustering(25). Amplitude differences between clusters may have skewed the analysis, so we performed L2 normalisation, which is a standard approach in clustering analyses. Spectral clustering was selected (over k-means etc) as detecting clusters on a manifold embedded in a higher dimensional parameter space, capturing the majority of the variation of the data in the space whilst reducing complexity. Succinctly, this permits us to find patterns without noise emerging from the high dimensionality of EEG data (as each lead is, in essence, a dimension of parameter space).

We tried values of k (number of clusters) from 2 to 10 and based on the silhouette score we selected an appropriate score, with the highest silhouette score selecting the best value of k. The Calinski-Harabasz score was also computed as a secondary metric. Stability was estimated using the adjusted Rand index (ARI) and split-half reliability.

We also performed clustering of the ECG data with simple k-means clustering (as ECG is much lower dimension than EEG), and explored how ECG cluster membership predicted CA cluster membership, to delineate how much of the variation was from cardiac behaviour, rather than head/neck conduction variation, using Cramer’s V to assess the membership predictions.

#### Comparing EEG response to the heartbeat with the cardiac artefact

We next compared heartbeat-locked scalp EEG between the heart-rate-matched control group (BM) and the isoelectric group (IE). For each participant, accepted EEG epochs time-locked to the R peak were averaged to produce a single participant-level sensor-by-time representation. The analysis therefore compared participant-level heartbeat-locked responses between groups.

The rationale for this comparison was that heartbeat-locked activity in the BM group (*EEG_BM_*) was expected to contain both a cortical heartbeat-evoked potential (HEP_BM_) and non-neural cardiac artefact (CA_BM_):

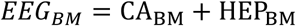

In contrast, heartbeat-locked activity in the IE group ( *EEG_IE_*) was assumed to contain predominantly cardiac artefact ( CA_IE_ ), with minimal or no scalp-detectable cortical HEP (HEP_IE,residual_)

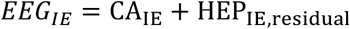

The observed group contrast (Δ_BM−IE_) can therefore be expressed conceptually as:

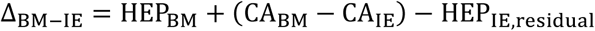

HEP_IE,residual_ is assumed to be close to zero by ensuring that there is no scalp recordable EEG in the IE group, and we attempt to match the CA profiles in both groups by matching heartrates. That is, the IE group therefore provides an empirical reference for the distribution of heartbeat-locked scalp activity in the presumed absence of a detectable cortical HEP. Sensor–time regions in which the BM group differed from this reference were interpreted as evidence for the spatiotemporal characteristics of the HEP.

To compare the two groups we used spatiotemporal cluster permutation with 5000(26), permutations. To ensure the analysis wasn’t affected by non-R-peak locked differences between the two groups, rigorous methods were adapted from Steinfath et al(27), in which non-heartbeat locked epochs were averaged and subtracted from the true HEP (Note this is the surrogate heartbeat method from their paper, not the pseudotrial method which does not apply to condition comparisoned HEP). We present both the cluster permutation with and without this surrogate heartbeat null.

Permutation tests are non-parametric and remain valid even in the context of asymmetric group sizes(26, 28), hence no subsampling or other methods were required to balance the groups. Spatiotemporal clustering was performed using calculated adjacency matrices for the 10-20 montage. Surrogate heartbeats were created (randomly) at a minimum of 0.25 seconds from the R-peak. As well as the surrogate heartbeat correction, we applied correction for nuisance effects, with Freedman-Lane methods(29, 30). The Freedman-Lane statistics were used to account for group differences.

We also compared the ECG waveforms between groups using an equivalent permutation procedure. This analysis assessed whether there were detectable group differences in the recorded ECG signal. The absence of a significant ECG difference was not interpreted as proof that all electrical and cardiomechanical components of cardiac artefact were equivalent between groups.

A baseline of -0.2 to 0 was used, reflecting common practice(23). Our group has previously used a smaller window(12, 31), but it was observed the QRS complexes in the IE group appeared wider, on average, than the BM group, and as such a narrow baseline from -0.2 to -0.04 would preferentially shift the IE group, introducing bias. The results of the -0.2 to -0.04 are included in supplementary materials – showing similar clusters to our primary analysis.

### Covariate-corrected spatiotemporal permutation

For each sensor and time point within the analysis window (50 to 900 milliseconds post-R-peak), the evoked amplitude was modelled as a function of Group membership, controlling for nuisance covariates. The design matrix, Z included:

- Heart Rate: Mean HR (z-scored).
- Global Amplitude Scale: A participant-level scalar quantity derived from the mean Global Field Power (GFP) within the analysis window (z-scored). This was included to control for global inter-subject variability in signal power, tissue conductivity or similar.

At each electrode and time point, we tested whether heartbeat-locked EEG amplitude differed between the BM and IE groups after accounting for the prespecified nuisance covariates. Because the nuisance covariates could be correlated with one another, their effects were modelled using ridge regression. The ridge penalty parameter, λ, was selected using 10-fold cross-validation.

Statistical significance was assessed using a Freedman–Lane permutation procedure(29). First, a reduced model containing only the nuisance covariates was fitted to the data. The residuals from this model represent variation that was not explained by the nuisance covariates. These residuals were permuted across participants 5,000 times and, for each permutation, added back to the values predicted by the nuisance model. These generated datasets represented the null hypothesis of no systematic group effect while preserving the effects of the nuisance covariates.

For the observed and each permuted dataset, the unique effect of group membership was calculated at every electrode and time point using a semi-partial-correlation-based t-statistic. This statistic quantified the association between group and EEG amplitude after accounting for the nuisance covariates.

To correct for testing across multiple electrodes and time points, samples exceeding a two-sided, pointwise threshold of p<0.05 were grouped into clusters based on spatial electrode adjacency and temporal continuity. The mass of each cluster was calculated as a sign-normalised sum of T values. For every permutation, the largest cluster mass was retained, producing a null distribution of maximum cluster masses. Each observed cluster was compared with this distribution to obtain a family-wise-error-corrected p-value.

### Sensitivity Analyses

Two exploratory sensitivity analyses were performed.

First, the primary comparison was repeated after applying a surface-Laplacian transform. This attenuates spatially broad, volume-conducted fields and reduces the dependence of scalp polarity on the average reference. It was used to test whether the frontocentral group difference persisted when the influence of distant voltage fields was reduced.

Second, the analysis was repeated after cardiac-like ICA components had been removed from both groups. This analysis tested whether the group difference persisted after conventional cardiac-artefact correction and whether ICA attenuated the putative HEP. This analysis used the average-reference data.

## RESULTS

100 records from 40 participants were available with isoelectric EEGs, where the EEG was of appropriate quality and duration. After excluding those with too few (usable) epochs, 39 were viable for further analysis. A dataset of 300 bpm-matched records was found, and after exclusion this retained 118 participants, which maintained the target participant ratio of 3:1. The included participants ages were 49.8(18.3) for the IE group, and 27.3(22.9) for the BM group. 48.7% participants were female in the IE group, compared to 57.6% in the BM group. Details of the included participants, including the medical and medication history available from the EEG reports, are shown in Table 1.

**Table 1.**
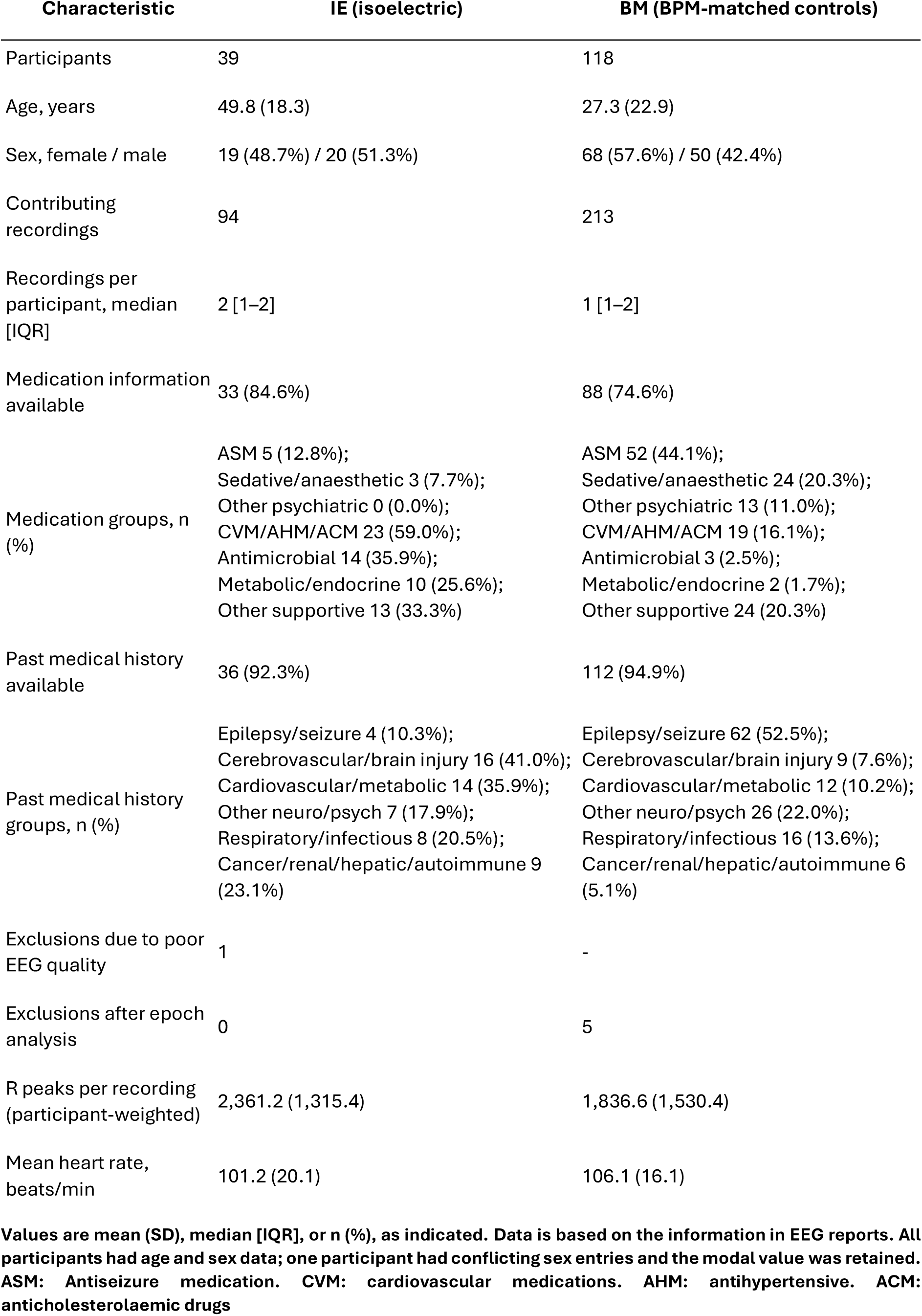
Participant characteristics and record inclusion.

### Morphological Analysis of the Cardiac Artefact and ECG

Spectral clustering of the IE group, focussing on the period from 0.1 to 0.9 seconds post R peak (in the time warped data), identified an optimal interindividual set of seven morphological clusters, according to the silhouette score (figure 1, top). This did not have good agreement with the Calinski-Harabasz score, which had a slope tending towards a trivial low-rank solution. Thus, support for precisely seven clusters was not consistent across the two selection indices (Figure 1).

**Figure 1:**
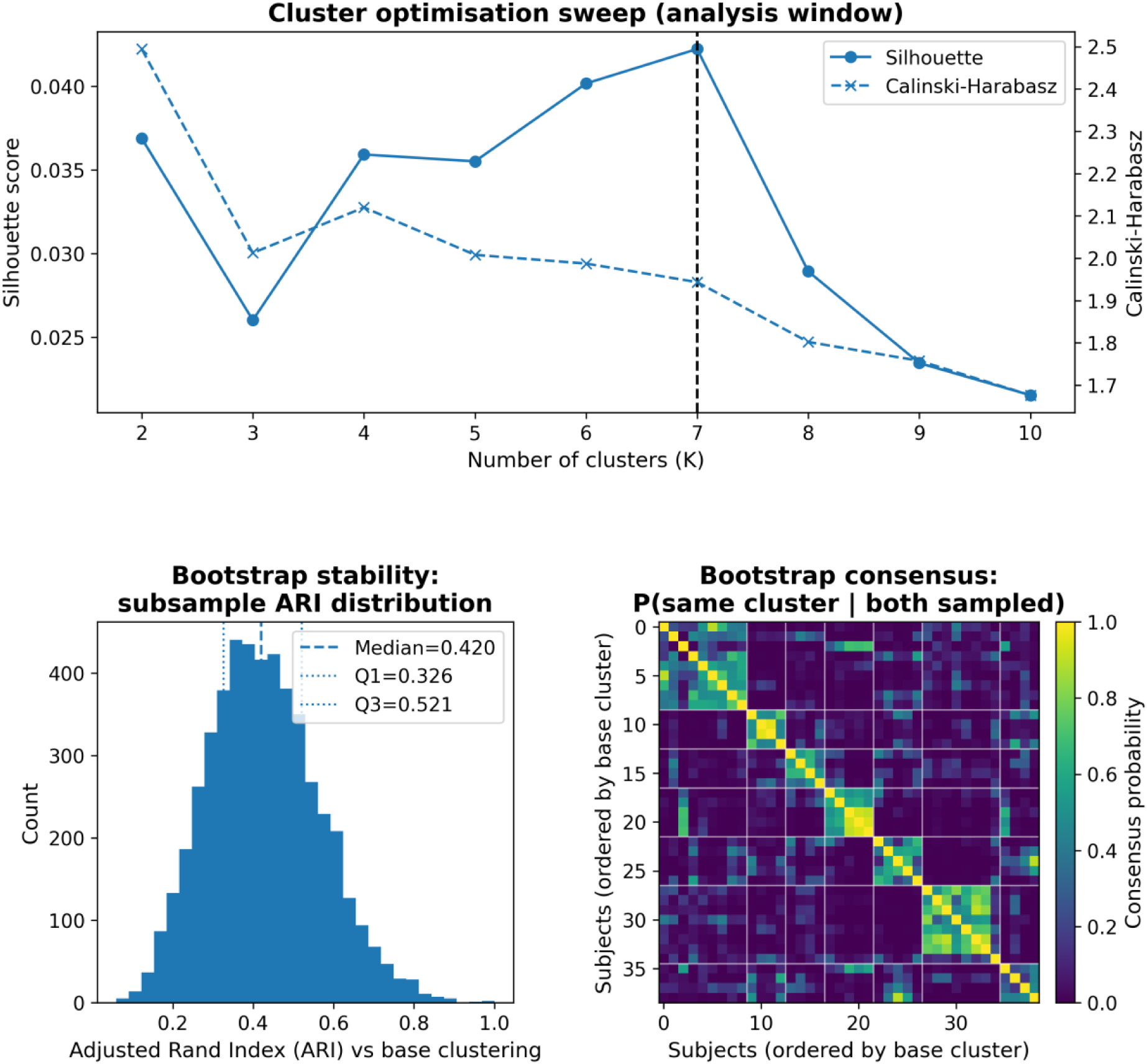
Top: Performance for varying cluster count (k) in the spectral clustering. The vertical dotted line indicates the selected value, based on the silhouette score. Note that the Calinski-Harabasz score did not agree well with the silhouette score, but the silhouette score appeared reliable with a clear peak: k=7**. Bottom Left: Distribution of the adjusted Rand index (ARI), for the cluster solution selected at k=7. Bottom Right: Cluster consensus grid for the k=7 cluster solution.**

Reproducibility and stability analysis identified a bootstrapped ARI median of 0.42 (figure 1, bottom left), and a median split-half reliability of 0.989, together these indicate a relatively stable membership between the clusters, at least within this limited sample size.

The clusters differed both in temporal (waveform) and topographical distribution (figure 2). Six of Seven clusters had a prominent posterior or left posterior negativity. Clusters zero and five had the most members. Cluster zero had a very focal negativity at the occiput, whereas cluster five had a broader field with an isopotential line from the left anterior to the right posterior. The remaining five clusters varied in terms of spatial extent and lateralisation, but all had a similar posterior negativity to cluster zero.

**Figure 2:**
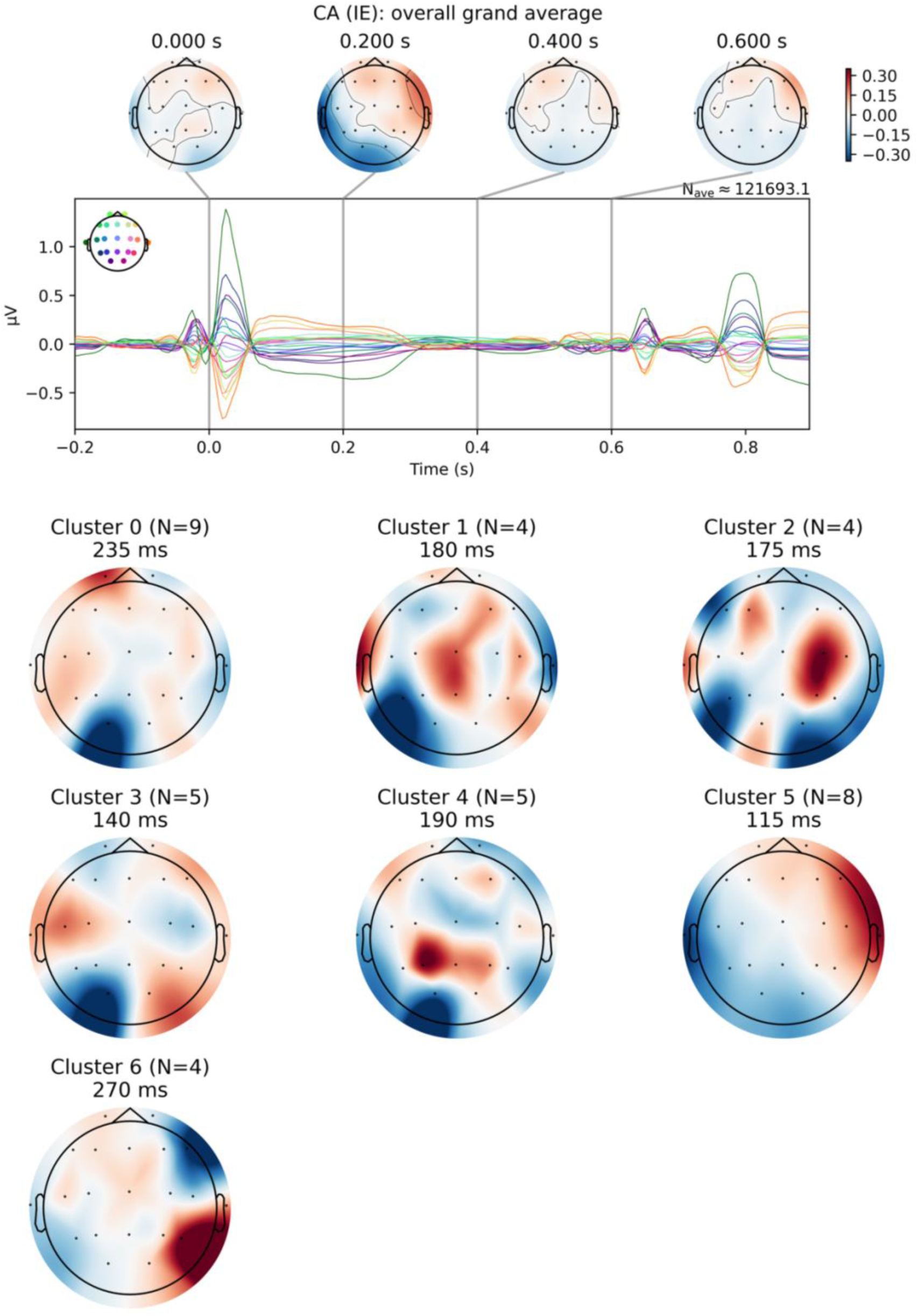
Top: global average for all isoelectric EEGs – showing the average cardiac artefact (CA) morphology. Bottom: Interindividual morphological clusters of the cardiac artefact, from 0.1 to 0.G seconds across the RR interval. N is the number of subjects in the isoelectric group which were best explained by a given cluster.

Hence, the present study found interindividual variation in the morphological characteristics of the CA – and it did not follow a simple pattern. K-means clustering of the ECG suggested an optimal silhouette score at k=2. These two clusters did not successfully predict the CA clusters (Cramer’s V: 0.50, p=0.14). Permutation analysis comparing the ECG between groups did not identify any significant differences (0 significant time points, minimum p=0.12).

### Evoked potential analysis – group comparison

The ECG waveforms did not differ significantly between the IE and BM groups in the permutation analysis. No significant temporal clusters were identified (minimum pointwise p=0.12). This analysis did not, however, test equivalence between the groups.

In the primary permutation analysis for mass-univariate comparison of the IE and BM base dataset (with preserved CA) demonstrated two significant clusters of difference between the evoked responses (Figure 3); in theory this represents differences attributable to the brains with cortical activity versus those without. The first was fronto-centro-temporal, scalp-negative cluster and spanned from 0.185 to 0.505 seconds (mass=506, p=0.009). The second was a left posterolateral, scalp-positive cluster and spanned from 0.15 to 0.31 seconds (mass=290, p=0.049). After correction with surrogate heartbeats, the first cluster persisted, but its temporal span reduced to 0.09 to 0.365 seconds (mass=414, p=0.016) and shifted to have a more right-sided emphasis. The second cluster persisted, with minimal change in its parameters (mass=314, p=0.03, timespan: 0.145 to 0.31 seconds).

**Figure 3:**
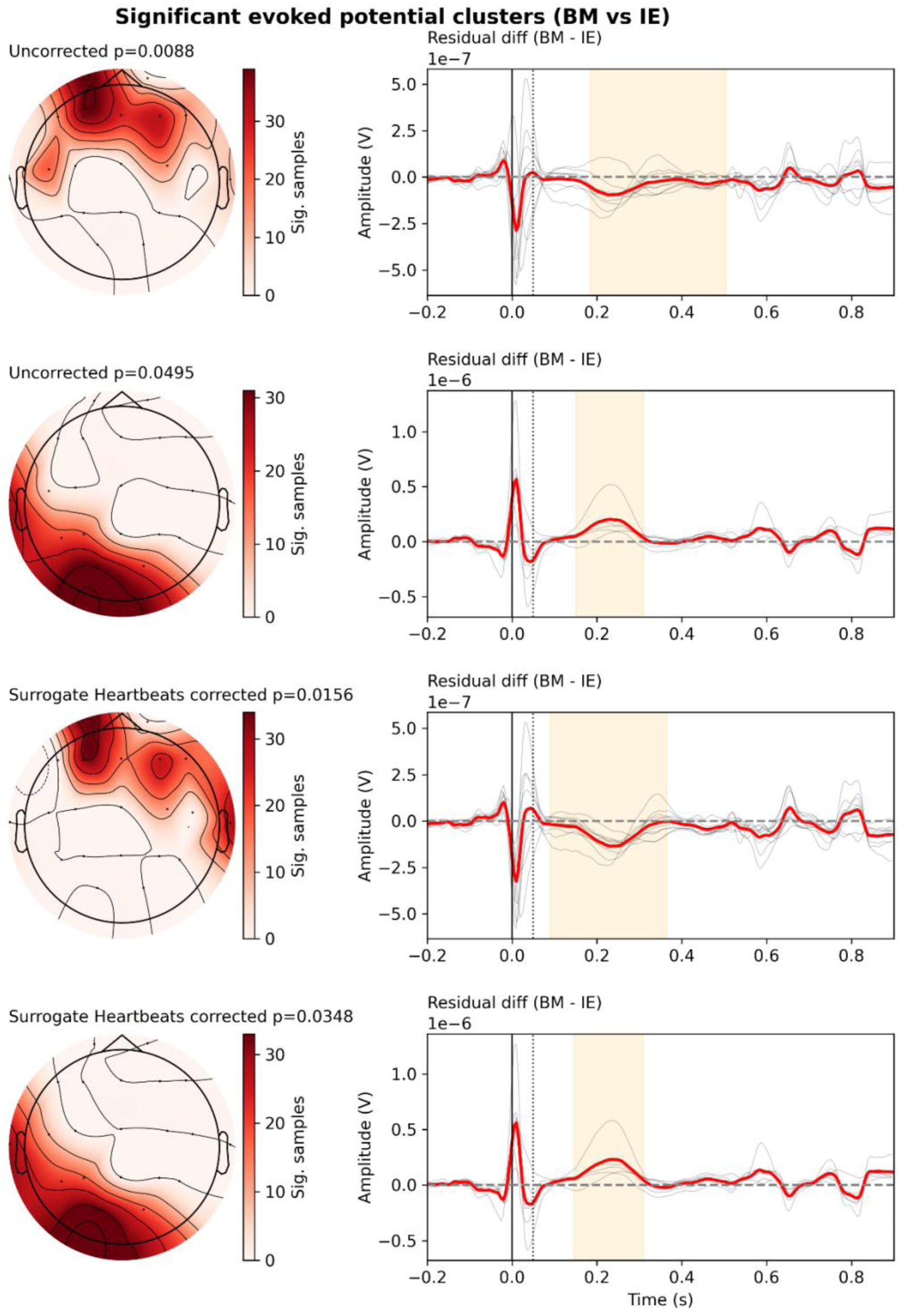
Significant differences in the heartbeat-evoked potential (HEP) compared to the cardiac artefact. This comparison is operationalised as the group difference between the heartrate-matched controls (BM) and the isoelectric group with isoelectric EEG (IE). We show here both clusters identified before (“uncorrected”) and after surrogate heartbeat correction.

### Sensitivity analysis

In the sensitivity analysis, repeating the permutation analysis with the Laplacian montage suppressed the posterior cluster, but the anterior negative cluster persisted (Figure 4). The distribution became more left-temporal, but fronto-central leads remained involved (before surrogate heartbeat correction: cluster mass=346, p=0.01, timespan: 0.175-0.405 seconds, after surrogate heartbeat correction: cluster mass=559, p=0.002, timespan: 0.175-0.505 seconds).

**Figure 4:**
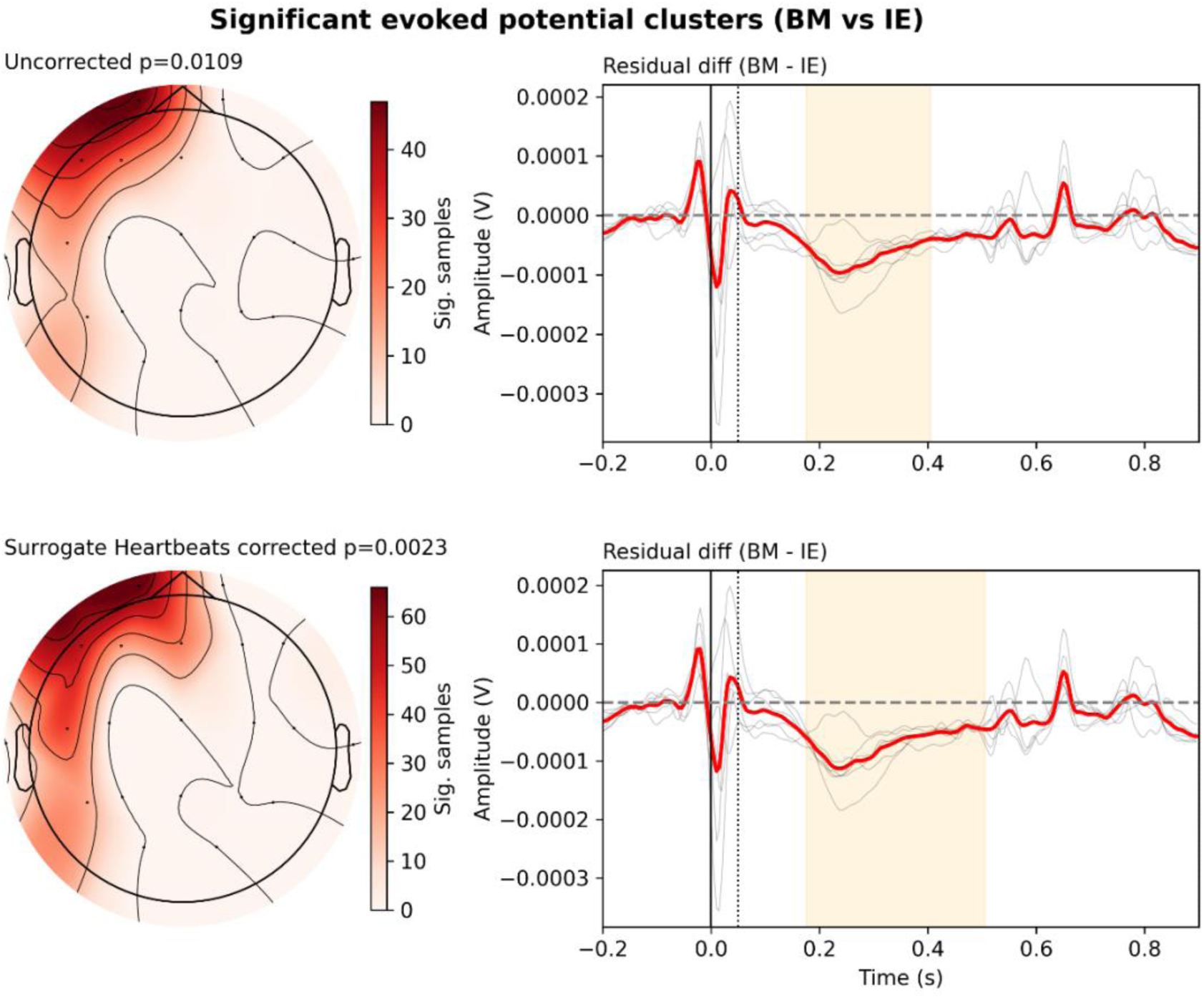
Significant differences in the heartbeat-evoked potential (HEP) compared to the cardiac artefact, with the Laplacian montage. Here, locally compared voltages demonstrate a significant cluster of negative difference in the heartrate-matched (BM) subjects, compared to the isoelectric EEGs (mass=55S, p=0.002, timespan: 0.175-0.505 seconds).

Finally, the group comparison was repeated after removal of manually identified cardiac-artefact-like ICA components from both groups (Table 2). Before surrogate-heartbeat correction, a negative frontocentral cluster was observed but did not reach statistical significance (cluster mass =287, p=0.059). After surrogate-heartbeat correction, a significant negative frontocentral cluster was identified from 170 to 385 ms (cluster mass =321, p=0.0487, Figure 5). Thus, a negative frontocentral BM–IE difference was observed across the average-reference, surface-Laplacian and ICA-corrected analyses, although in the ICA-corrected data it reached statistical significance only after surrogate-heartbeat correction.

**Figure 5:**
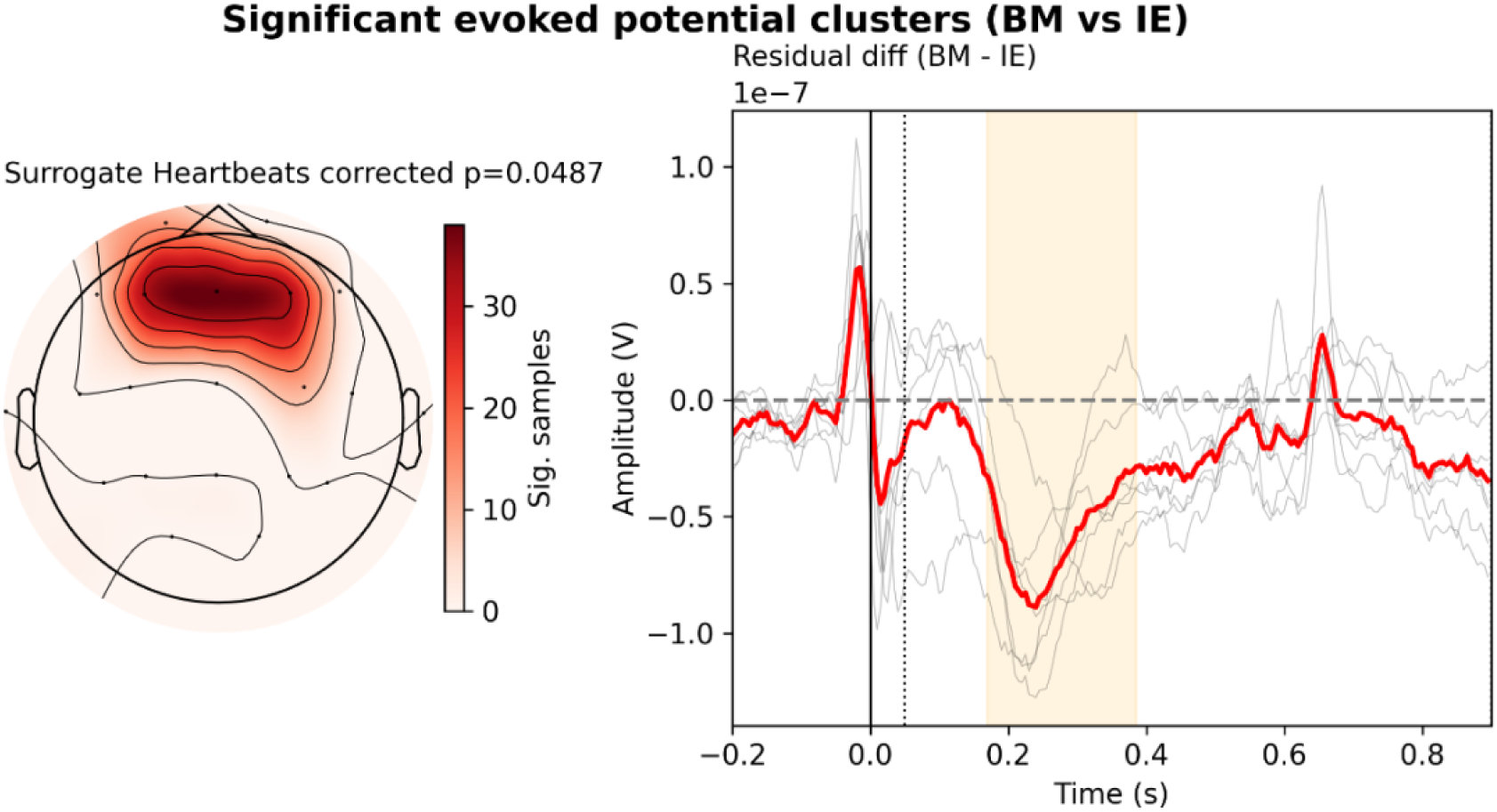
Significant differences in the heartbeat-evoked potential (HEP) compared to the cardiac artefact, following ICA to remove cardiac artefact. With removal of the cardiac artefact, a single frontocentral cluster emerges after surrogate heartbeat correction. Prior to surrogate heartbeat correction, there was no significant cluster (mass=287, p=0.05S), but the cluster becomes significant after correction (mass=321, p=0.04S, timespan: 0.17 to 0.385 seconds). As with other montages and analyses, it is a negative difference between the HEP and the isoelectric baseline.

**Table 2.** Signal removed by ECG-informed ICA.

| ICA removal characteristic | IE (isoelectric) | BM (BPM-matched controls) |
| --- | --- | --- |
| Participants with $\geq 1$ component appeared to be cardiac artefact | 32 (82.1%) | 73 (61.9%) |
| Rejected components, total | 117 | 238 |
| Rejected components per participant, median [IQR] | 3.0 [1.0–5.0] | 1.0 [0.0–3.0] |
| Rejected share of fitted components, median [IQR] | 14.3 [5.0–23.8]% | 9.5 [0.0–16.2]% |
| Participants with 0 / 1 / 2–4 / $\geq 5$ rejected components | 7 (17.9%) / 4 (10.3%) / 17 (43.6%) / 11 (28.2%) | 45 (38.1%) / 19 (16.1%) / 38 (32.2%) / 16 (13.6%) |
| Rejected-component QRS:post-T energy ratio, median [IQR] | 2.9 [2.4–4.3] | 2.6 [2.2–3.6] |
| Rejected-component QRS peak-to-peak, a.u., median [IQR] | 0.8 [0.3–2.6] | 0.4 [0.2–0.8] |

**Details relating to the independent component analysis (ICA) based removal of the cardiac field artefact, with manual identification. The relative proportion of the QRS-complex period to the post-T period (region with minimal ECG artefact) is used as a heuristic of the specificity of the components selected to cardiac artefact.**

## DISCUSSION

The present study demonstrates the morphological characteristics of the CA in the absence of the heartbeat-evoked potential. Previous HEP studies in the brain-dead population(32) have not compared groups with and without brain activity. The present study presents evidence delineating, at the scalp, the neural response to the heart from heartbeat-associated artefact and describing the morphologies of the HEP and the CA as separate entities.

### The Morphology of the Cardiac Artefact

The CA was spatially broad, consistent with it being a distant, high amplitude source distributed across the overlying tissues (Figure 2). The topographic distribution will likely overlap with and (therefore possibly distort) the HEP.

Interindividual breakdown of the epoched data in the IE group indicated seven groups (Figure 1, top). The variability of the CA in the population is unknown, but in this relatively small sample, we can see significant variability of the spatiotemporal distribution of the artefact, but with typically asymmetrical distributions of the artefact in the T wave window. We note that many clusters are strongly focussed on the occiput and may well represent pulsatile movement artefact against the pillow/bed – as the IE group were (likely) all supine, though this is post-hoc speculation.

While the clusters varied in morphology, the cluster solution itself was stable (an average adjusted Rand index of 0.42 indicates a moderate level of agreement across bootstrapped subsamples (Figure 1, bottom left). It is impossible to interpret whether some of the clusters of variation represent interindividual subject position (such as head rotation) or inherent interindividual differences such as tissue conductivity and thoracic or cephalic geometry.

The failure of the ECG clustering (in this small sample) to predict the scalp behaviour can be explained by one or a combination of two phenomena. Firstly, the individual soft tissue conductivity between individuals is highly variable and linked to body habitus and height. Secondly, the CA has an unknown proportion caused by the CBA. The former is well established, and evidenced by the head-turn manipulations by Dirlich et al(5). The latter is unquantifiable by current methods, and is typically overlooked in analyses.

### The Morphology of the Heartbeat Evoked Potential

The evoked response significantly differed from the averaged CA (figure 3). A frontocentral cluster (similar to the expected frontal component of the HEP in the literature), and a posterior cluster were identified. The initial (uncorrected) frontocentral HEP-like cluster spanned a broad timebase, including both the early and late HEP periods. However, after surrogate heartbeat correction, the span contracted to the early HEP period – suggesting that the latter part of the cluster represented non-heartbeat-locked drift. This does not necessarily mean that the late HEP found in other studies is artefactual – it may well be an effect of the high heartrate in the studied population introducing obfuscating noise in the late epoch (which is visible from the oscillatory nature of the T statistic in this part of the plot on figure 3). Hence, we cannot make claims about the nature or morphology of the late HEP in this analysis. The posterior may be group differences in CA, but also may represent the positive field of a dipolar HEP.

Montage manipulation supported an analysis unbiased by montage effects, allowing a more holistic consideration of the voltage effects. The span of the frontocentral HEP-like cluster extended further into the cardiac window when a Laplacian montage was used, and this persisted after surrogate heartbeat correction. The Laplacian, while being less able to detect broad scalp potentials, did not support the presence of a significant posterior difference between the groups. This is reassuring that the clusters seen were not an artefact of the average montage.

After ICA, the frontocentral HEP-like cluster became much less prominent and was obscured to the extent that it only emerged after surrogate heartbeat correction. This demonstrates firstly that the surrogate heartbeat method proposed by Steinfath et al can unmask genuine significant clusters (rather than being a purely conservative procedure)(27). Secondly, specifically in the case of this study, the emergence of the HEP after the surrogate-trial correction in the ICA-corrected data shows that HEP is susceptible to obfuscation under non-heartbeat-locked random fluctuations. This is explicable by its being such a low-amplitude potential, compared to many others. However, it is important to note that there were groupwise differences in the cardiac artefact identification between the two groups (Table 2).

The frontocentral morphology overlaps well with the range of leads targeted in other literature(33). This continues to support typical suggestions for generators. However, in the sparse electrode array available in this study, no source localisation is possible. The loss of the second part of the HEP (the late HEP) after surrogate heartbeat correction may be a reflection of greater artefact jitter in the later part of the epoch, due to subsequent heartbeats (though as noted above, it persists in the Laplacian analysis). This is something which likely cannot be completely controlled for statistically with our Freedman-Lane model (or similar mixed modelling) since interindividual heartrate variation is likely collinear with interindividual morphological differences.

### The Polarity of the HEP

Permutation analysis of the CA and HEP against the CA background permitted an unbiased estimation of the HEP’s polarity, suggesting a negative early HEP. This aligns with recent work by Gautier et al (34), who propose an early scalp-negative deflection of the HEP, though this relied on an assumption that ICA would remove the CA without excessively affecting the HEP. We did not find a late HEP cluster so cannot confirm or contradict their finding of a positive late HEP (after ∼300ms). The Laplacian montage avoids the theoretical leak from a posterior positive cluster into an anterior negative cluster and taken together with the average montage we can conclude with a degree of certainty that the difference in the residualised heartbeat-locked EEG between the BM and IE group was supportive of a negatively polarised HEP (at the scalp) and that this is not an artefact of montage choice. Studies using other montages (such as the Laplacian, or bipolar montages) have not been identified in a recent review(23) – though some studies fail to report the reference system used.

García-Cordero et al. showed that an interoceptive condition produced a significantly more negative frontally distributed HEP, while a paired intracranial study demonstrated increased broadband activity within canonical interoceptive regions, including the posterior insula, inferior frontal gyrus, amygdala and somatosensory cortex(35). Aligning the two parts of their studies, if the HEP is related to signals in the structures studied stereotactically, then this suggests the heartbeat associated increase in broadband activity in these regions projects a scalp-negative EP (between 170 and 400ms post R peak) under an average montage. This aligns with the present study.

In intraindividual analysis, once the cardiac-artefact background is accounted for, the early HEP is expressed as a negative frontocentral scalp field. Within a fixed montage and sensor–time region, greater component expression should therefore appear as a more negative voltage, whereas attenuation should appear as a shift towards zero. In interindividual analysis, however, the high variability in the CA (and the fact that its overlap may well confound temporal ICA methods to an extent variable between participants) means that inferences in the HEP amplitude need to be made with caution.

The early HEP may include ascending cardiac-afferent activity, prediction-error-related activity, precision or gain modulation, recurrent integration, and heartbeat-triggered coordination of ongoing cortical dynamics. Establishing its scalp polarity permits the direction of component modulation to be interpreted, but does not alone identify which of these computations generated it.

### Artefact correction and accurate HEP recovery

Analysis of the HEP in the purely amplitude domain (equivalent to a time-locked, phase-locked analysis, collapsed across frequencies) requires robust accounting for the CA, as well as other random effects. Attempting to remove the CA with ICA seemed to suppress the HEP. While this might not be generalisable to higher dimension analyses (with higher lead counts and therefore higher component counts), this may imply that ICA for CA removal carries the risk of affecting (if not completely removing) the HEP. Even with a higher component count, the fundamental temporal collinearity between the HEP and the CA may make them intrinsically overlap in ICA solutions.

As it stands, the present study suggests that extreme care need be taken when using ICA in this way, and it is hard to know how researchers can realistically infer the extent to which ICA is affecting their HEP, and it would be challenging to know for certain that ICA had not differentially affected the HEP between participants or records.

The ECG did not successfully predict the CA cluster. This may be a lack of power due to the small sample size (39 participants). However, using the ECG to estimate the CA may implicitly fail on two counts. First, the ECG morphology may not predict the scalp artefact if most of the variation depends on anatomy of the torso and neck, rather than cardiac behaviour. Secondly, the ECG will only predict the CFA, but not the CBA. To successfully remove CA from the HEP, approaches might orthogonalize the signals in a mathematically sound manner, rather using assumptive heuristics. Until methods for this are well established, interindividual comparisons, or within-individual studies in which cardiac state is modulated pose a high risk of error.

ICA has been shown by Buot et al to successfully remove the beat-to beat fluctuation in the heartbeat-evoked response, when carefully aligned with the cardiac behaviour(36). This improves recovery of relevant signal when the focus of the analysis is directed at modulations of the HEP which do not change in alignment with cardiac behaviour, but one would lose potentially relevant information about the beat-by beat processing of a variable cardiac signal. As such, ICA poses a risk in HEP studies proportional to the correlation of the independent variable and cardiac behaviour.

Traditionally, artefact correction has been attempted with ICA or similar methods, focus on temporal windows with significant CA (e.g. before or after the T wave)(23), or within participant comparison to account for interindividual differences in the CA(12). The current findings demonstrate the potential pitfalls in artefact correction methods, but within-participant methods remain feasible.

### Limitations

Intracranial injury and skull or scalp abnormalities may have altered volume conduction in some IE participants. The available clinical metadata did not permit these effects to be quantified. Electrocorticography records amplitudes of the CA less than a quarter of the amplitude at the scalp(37), and that is likely an underestimate of the difference as electrocorticographic studies implicitly have an overlying breach effect. Furthermore, there is a skull-brain conductivity ratio of around 25±7(38). Hence, it is unlikely that any extent of brain tissue conductivity changes would have significantly modified the CA as recorded at the scalp at the group level.

There are several limitations associated with the methodology of this retrospective study. Firstly, this study was designed based on the assumption that all evoked potentials would become absent in a sufficiently damaged brain, as indicated by isoelectricity. There is some evidence that, while scalp isoelectricity reflects an absence of cortical EEG activity, deeper structures may still generate EEG signals(39) (though this was not demonstrated in brain death). As the HEP appears cortically generated, it is unlikely to be preserved even if deeper activity (such as brainstem auditory evoked potentials) persists(32), and this is supported by previous analysis of the HEP in brain death(32). Also, as noted above, we only require that there is a systematic attenuation of brain activity to the point of it not being scalp-detectable for the HEP to be functionally zero for the purposes of this study.

Another limitation is the use of reportedly normal EEGs from a clinical dataset rather than true healthy controls. This was done to create an equivalent dataset of normal EEG matched to the brain death (rather than using a dataset of known healthy controls but with possible differences in the recording set up). An EEG reported as normal may not preclude brain abnormalities, and as the HEP is not considered in a normally-reported EEG, the HEPs included in the controls may not be “normal.” Groups differed in age, medications and medical history. Finally, the groups may have systematically differed in body habitus or in non-modelled cardiodynamic behaviour such as stroke volume, which could have introduced uncorrected systematic differences in CA.

## CONCLUSION

The present study empirically explores the CA and the HEP as distinct entities using a real-world dataset. In this naturalistic clinical sample, we demonstrate significant variation in the morphology of the CA, which complicates interpretation of artefact correction methods. We find evidence supporting the notion that the HEP is a distinct entity from the underlying CA. Finally, we demonstrate that in the studied sample, the HEP has a frontocentral distribution and that the (early) HEP is negative at the scalp.

## Large Language Model use declaration

ChatGPT (Open AI) was used to improve code efficiency, though all methods were originally written by RK, or from analytical packages which are already peer reviewed and cited.

## Data Availability Statement

All raw data is sourced from the Temple University EEG Corpus(15), with their permission. All code is available in the HEPPy project on GitHub: ([link](https://github.com/ClinicalAffectiveNeuroscienceLab/HEPPy)). Detailed preprocessing logs are also available.

## SUPPLEMENTARY MATERIALS

### HEP cluster permutation analysis with a narrower baseline

Given the discrepancy between the selected baseline in this analysis (which was chosen based on noted differences between the groups in terms of the QRS width) a subsidiary analysis is attached here using the baseline used elsewhere by this group (-0.2 to -0.04).

**Figure S1:**
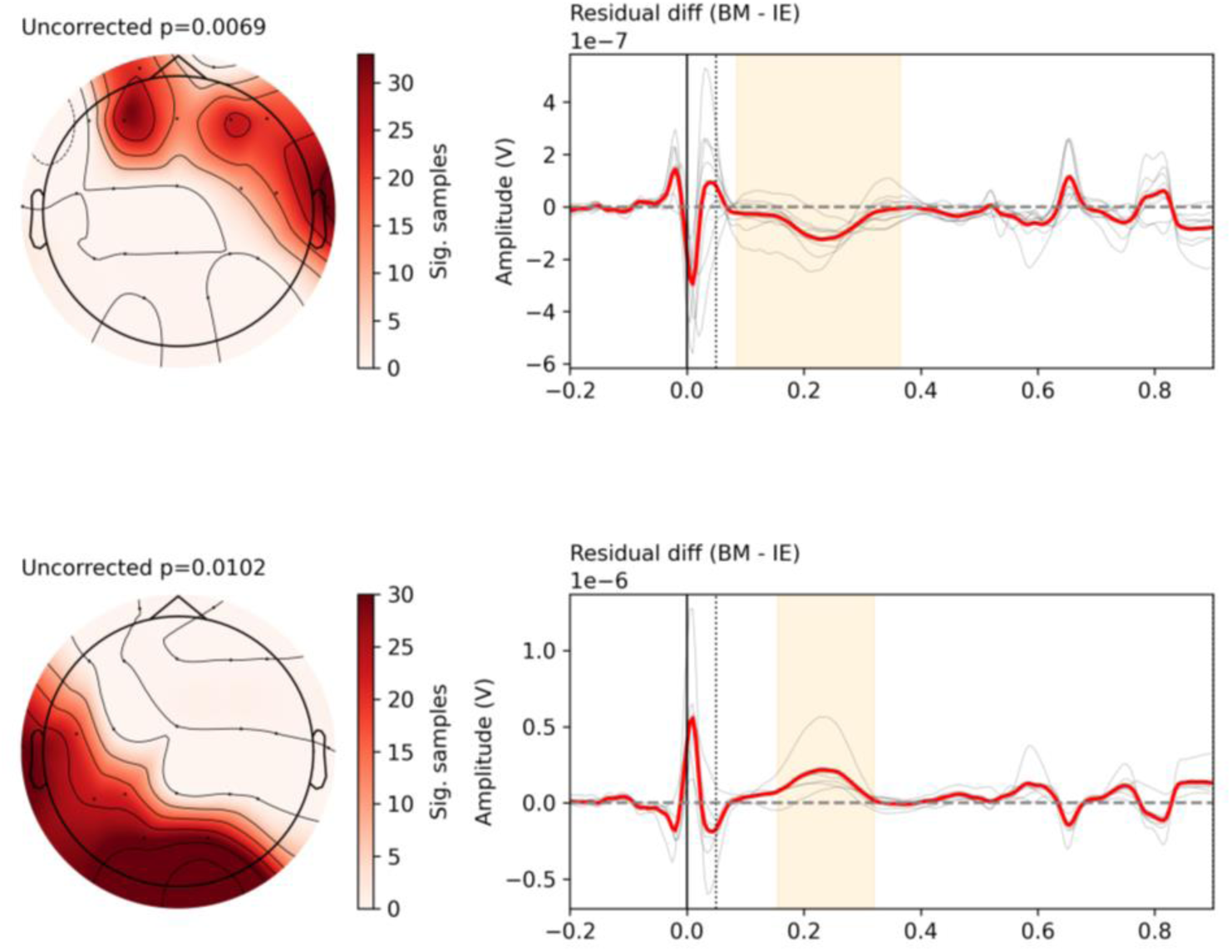
Average montage significant clusters comparing the isoelectric and control groups when using a baseline of -0.2 to -0.04. BM: heartrate matched. IE: Isoelectric.

